# The environment-sensing aryl-hydrocarbon receptor inhibits the chondrogenic fate of modulated smooth muscle cells in atherosclerotic lesions

**DOI:** 10.1101/2020.04.21.049890

**Authors:** Juyong Brian Kim, Quanyi Zhao, Trieu Nguyen, Milos Pjanic, Paul Cheng, Robert Wirka, Manabu Nagao, Ramendra Kundu, Thomas Quertermous

## Abstract

**Introduction:** Smooth muscle cells (SMC) play a critical role in atherosclerosis. The Aryl hydrocarbon receptor (AHR) is an environment-sensing transcription factor that contributes to vascular development, and has been implicated in coronary artery disease (CAD) risk. We hypothesized that AHR can affect atherosclerosis by regulating phenotypic modulation of SMC.

**Methods:** We combined RNA-Seq, ChIP-Seq, ATAC-Seq and in-vitro assays in human coronary artery SMC (HCASMC), with single-cell RNA-Seq (scRNA-Seq), histology, and RNAscope in an SMC-specific lineage-tracing *Ahr* knockout mouse model of atherosclerosis to better understand the role of *AHR* in vascular disease.

**Results:** Genomic studies coupled with functional assays in cultured HCASMC revealed that *AHR* modulates HCASMC phenotype and suppresses ossification in these cells. Lineage tracing and activity tracing studies in the mouse aortic sinus showed that the *Ahr* pathway is active in modulated SMC in the atherosclerotic lesion cap. Furthermore, scRNA-Seq studies of the SMC-specific Ahr knockout mice showed a significant increase in the proportion of modulated SMC expressing chondrocyte markers such as *Col2a1* and *Alpl*, which localized to the lesion neointima. These cells, which we term “chondromyocytes” (CMC), were also identified in the neointima of human coronary arteries. In histological analyses, these changes manifested as larger lesion size, increased lineage-traced SMC participation in the lesion, decreased lineage-traced SMC in the lesion cap, and increased alkaline phosphatase activity in lesions in the *Ahr* knockout compared to wild-type mice. We propose that *AHR* is likely protective based on these data and inference from human genetic analyses.

**Conclusion:** Overall, we conclude that *AHR* promotes maintenance of lesion cap integrity and diminishes the disease related SMC-to-CMC transition in atherosclerotic tissues.

## Introduction

Coronary artery disease (CAD) is driven by both genetic and environmental risk. The contribution of genes and the environment is estimated to be ~50% each based on large human genetic studies.^1,2^ Furthermore, genes can modify the disease process promoted by environmental exposures.^3^ Such interplay between genes and the environment has been difficult to study due to lack of power and lack of well-documented individual level measurement of exposure dosage.

The aryl hydrocarbon receptor (AHR) is a ligand activated transcription factor that is induced by both endogenous and exogenous compounds including environmental pollutants such as dioxin, polycyclic aromatic hydrocarbons (PAH), tobacco smoke, and organic waste products which have been linked to cardiovascular mortality.^4,5^ AHR has established roles in the activation of detoxifying enzymes in response to pollutant exposure, but also emerging roles in vascular development and homeostasis. Specifically, disruption of the *AHR* pathway resulted in reduced coronary artery formation, and increased cardiovascular birth defects in animal models.^6–8^ It is yet unclear whether AHR plays a protective role in the vasculature in response to environmental toxins or if AHR is the driver pathway that is responsible for pathological changes in the context of exposure to these atherogenic toxins.

In our previous work, we have shown that *TCF21*, a highly replicated gene associated with CAD, interacts closely with *AHR*.^9^ We also found that *TCF21* plays a central role in regulating SMC phenotype and in modulating SMC phenotypic modulation during atherosclerosis.^10–12^ However, the role of the AHR pathway in regulating SMC phenotypic modulation in atherosclerotic disease is not well understood.

In this study, we sought to further investigate the specific effects of *AHR* on the vascular smooth muscle cell phenotype in atherosclerotic disease. We show that *AHR* regulates the SMC phenotype by globally affecting the chromatin accessibility landscape of SMC and thus impacting expression of genes regulating development, cell migration, proliferation, as well as extracellular matrix organization and endochondral ossification. We further characterize the effect of *AHR* in a SMC-specific *AHR* knockout mouse model of atherosclerosis using single-cell sequencing and histological analysis, and conclude that *AHR* is a critical regulator of SMC contribution to atherosclerotic lesion.

## Results

### *AHR* modulation in HCASMC results in transcriptional changes relevant to vascular function and disease

In order to examine the transcriptional pathways regulated by *AHR*, we treated human coronary artery smooth muscle cells (HCASMC) with *AHR* siRNA or control siRNA and performed bulk RNA sequencing (RNA-Seq). Overall, at an adjusted p-value less than 0.00001 and fold change greater than 1.3, there were 1004 up-regulated genes, and 814 down-regulated genes (Figure 1a, Supplemental Table 1). The differentially regulated genes were enriched in curated biological processes including vascular development, response to organic substance, and regulation of cell migration (Figure 1b, Supplemental Table 1). To further validate these findings, we also transduced *AHR* overexpressing lentivirus into HCASMC and performed RNA-seq (Figure 1a, Supplemental Table 1). At an adjusted p-value less than 0.05 and fold change greater than 1.3, there were 151 up-regulated genes, and 327 down-regulated genes. We again found that vascular development, and response to chemical stimulus were among the top enriched pathways, in addition to cell cycle/cell division (Figure 1b, Supplemental Table 1).

**Figure 1.**
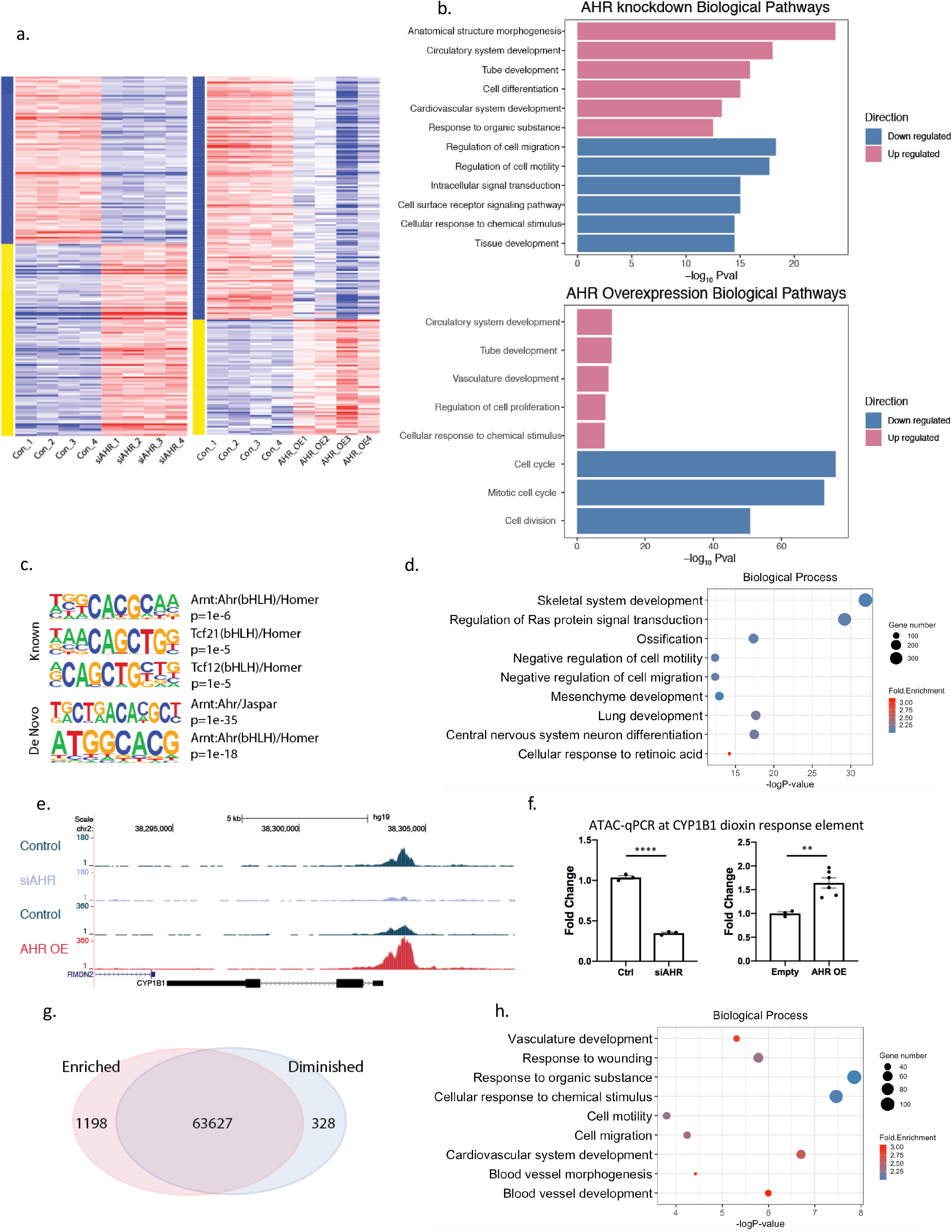
*AHR* modulates pathways relevant to vascular disease and induces chromatin accessibility in HCASMC. (a) Heatmap of differentially expressed genes in *AHR* knockdown (left), *AHR* overexpression (right) RNA-Seq data in HCASMC show a significant impact of *AHR* modulation on the transcriptome. (b) Top enriched biological pathways from differentially expressed genes of *AHR* knockdown (top), and *AHR* overexpression (bottom). (c) AHR ChIP-Seq peaks localize to AHR and TCF21 motifs with HOMER *known* and *de novo* analyses. (d) DAVID Gene Ontology analysis of AHR target genes identified by GREAT. (e) An ATAC-seq peak is visualized at the promoter region of *CYP1B1*, where the peak is diminished with knockdown and increased with *AHR* overexpression and (f) was confirmed with ATAC-qPCR (AHR knockdown by siRNA p<0.0001 (*left*), AHR overexpression p=0.0048 (*right*)). (g) *AHR* overexpression leads to an overall increase in open-chromatin regions. (h) DAVID Gene Ontology analysis of genes in the regions of increased chromatin accessibility signatures from *AHR* overexpression.

To determine which of the identified differentially expressed genes are directly regulated by AHR, we investigated the global pattern of AHR protein binding to chromatin by performing ChIP-Seq in HCASMC. The binding of AHR to the known *CYP1B1* dioxin response element (DRE) binding motif was confirmed as a positive control (Supplemental Figure 1). We identified a total of ~17000 peaks with the AHR ChIP-Seq study in this cell type. AHR and TCF21 motifs were both significantly enriched within the peaks (Figure 1c, Supplemental table 2). Genes identified by GREAT^13^ at AHR target loci were found by ontology analysis to be associated with terms related to development, ossification, response to retinoic acid, and cell migration (Figure 1d).

In addition to direct transcriptional regulation, we investigated whether AHR can affect gene expression in target loci through altering chromatin states. To examine this possibility, we performed the assay for transposase accessible chromatin (ATAC) followed by sequencing in HCASMC (ATAC-Seq). We found that *AHR* overexpression increased the chromatin accessibility in the promoter region of CYP1B1, and decreased accessibility when *AHR* was knocked down, as confirmed with ATAC-qPCR (Figure 1e, 1f). We then compared each treatment group to look for differentially enriched peaks. We found that *AHR* modulation had a significant impact on the global landscape of chromatin accessibility. When *AHR* was overexpressed, there were 1198 up-regulated peaks, and 328 down-regulated peaks at FDR<0.05 (Figure 1g). Globally, the open chromatin regions that enriched with *AHR* overexpression were clustered near genes related to response to organic substance, cardiovascular development and cellular migration and motility, mirroring those of the AHR ChIP-Seq enrichment results (Figure 1h, Supplemental Table 2).

### The *AHR* downstream pathway overlaps with the *TCF21* pathway and is enriched in regions of open chromatin in HCASMC

*TCF21* is a causal CAD associated gene that has been replicated in multiple racial ethnic groups and is activated in the setting of vascular stress to promote modulation of SMC to develop a fibroblast-like phenotype, i.e., as a fibromyocyte (FMC).^12^ To extend our previous studies that indicated a regulatory interaction between *AHR* and *TCF21*,^9^ we compared their downstream pathways as determined by combined RNA-seq and ChIP-Seq analyses. There was significant overlap in both up- and down-regulated genes with *AHR* and *TCF21* knockdown (Figure 2a). The pathways that were co-regulated at the mRNA level in these experiments were enriched for regulation of cell migration, development, ECM organization, and cell adhesion by GO analysis with DAVID (Figure 2b). There was also enrichment for cytokine-cytokine receptor interaction, TGF-β pathway and ECM-receptor interaction KEGG pathways (Figure 2c, Supplemental Table 3). We also considered the genes that were specifically regulated by *AHR* but not by *TCF21*. These genes were enriched for GO terms similar to the those identified for the overlap gene list including migration, but also specifically identified ossification/skeletal system development, collagen fibril organization, and SMC proliferation (Figure 2d, Supplemental Table 3).

**Figure 2.**
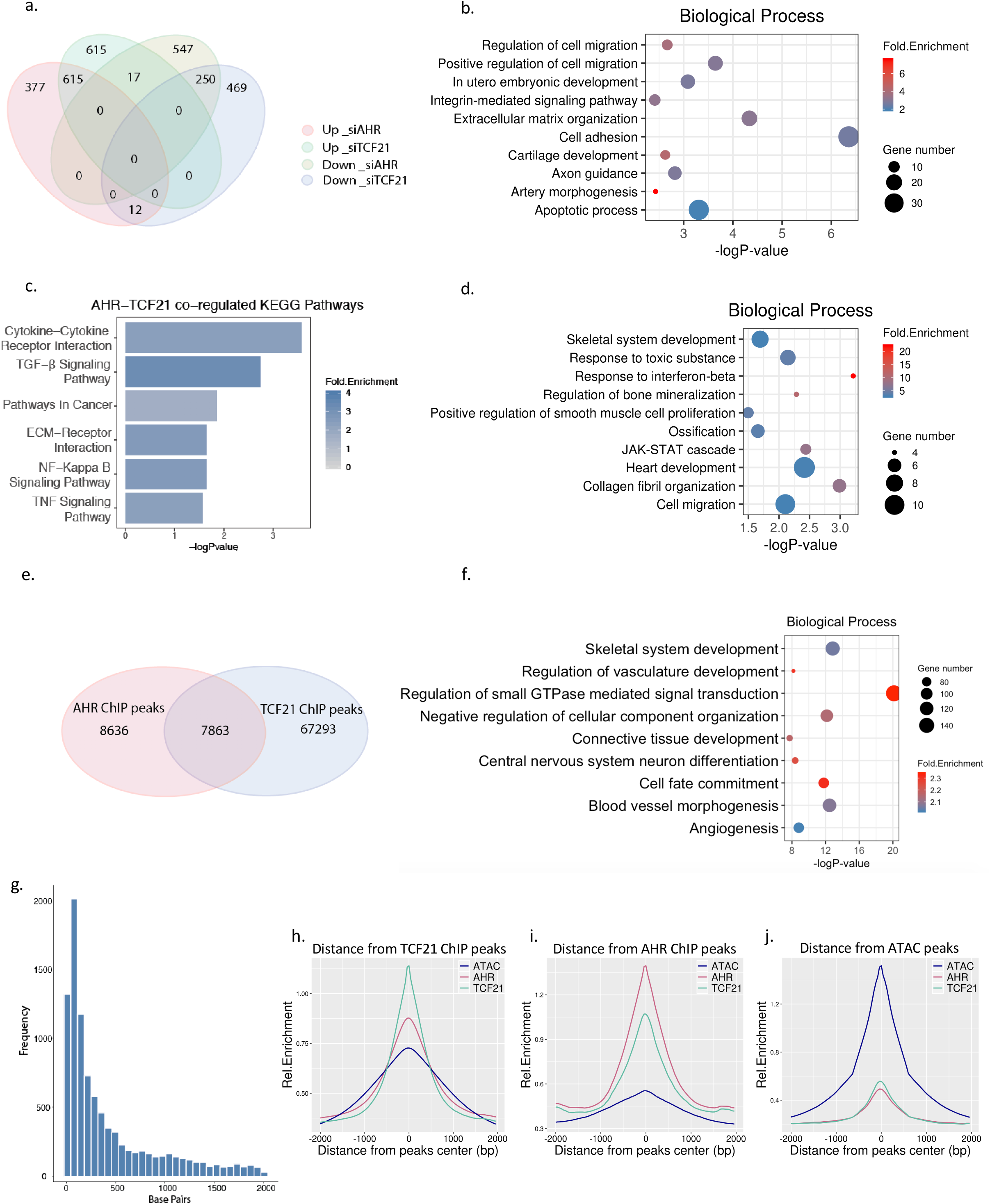
The *AHR* pathway overlaps *TCF21* downstream pathways and is enriched in open chromatin regions in HCASMC. (a) Comparison of *AHR* knockdown RNA-Seq and *TCF21* knockdown RNA-Seq results from HCASMC. A total of 1004 genes were up-reguated and 814 genes down-regulated in *AHR* KD, 1247 genes up-regulated, and 731 genes down-regulated in *TCF21* KD (FDR <0.00001, Fold Change >1.3). Among these 615 up-regulated and 250 down-regulated genes overlapped between the *AHR* KD and *TCF21* KD groups, respectively (Hypergeometric test p =1e-4642) (b) Top enriched biological pathways from up-regulated genes that are common to *AHR* KD and *TCF21* KD (c) The top KEGG pathways enriched from genes regulated by both *TCF21* and *AHR*. (d) Top enriched biological pathways unique to *AHR* KD. (e) AHR ChIP-Seq and TCF21 ChIP-Seq target loci share a significant number of intersected peaks. (f) DAVID Gene Ontology analysis of overlapping AHR and TCF21 binding peak genes identified by GREAT. (g) The distance between the AHR and TCF21 binding sites are on average 50-100 bp apart. (h-j) The peaks of TCF21 ChIP-Seq (green line), AHR ChIP-Seq (red line) and *AHR* overexpression ATAC-seq peaks (blue line) co-localize genome-wide, as shown by overlapping relative enrichment centered around each respective coordinates.

We then compared the AHR and TCF21 ChIP-Seq peaks to see which of these gene expression similarities were driven directly by joint binding at target loci. There was a significant overlap in the common peaks as identified by the two ChIP-seq studies (Figure 2e, p =1e-4642, hypergeometric test), with 48% of the AHR peaks intersecting with TCF21 target loci. The common peaks were enriched for skeletal system development, cell motility and migration, and development (Figure 2f, Supplemental Table 3). The distance between the summits of the peaks were compared, and the most common overlap was seen within 50-100 bp, but extended up to 1000 bp, suggesting that these two factors may interact through regulation of the local epigenome that mediates binding of AHR and TCF21 at these sites (Figure 2g). We further analyzed the chromatin accessibility at these ChIP-Seq peaks and found that AHR binding and TCF21 binding sites globally co-localized with open chromatin regions in HCASMC, suggesting that AHR and TCF21 affect the binding and function of each other (Figure 2h-j). Although co-immunoprecipitation studies failed to identify a direct interaction of these two transcription factors in solution (Supplemental Figure 2a), we found evidence of interaction when these proteins were cross-linked to chromatin by performing ChIP followed by western blotting, further supporting the hypothesis that the interaction of AHR and TCF21 is at least partially based on their joint chromatin scaffolding (Supplemental Figure 2b).

### Single-cell RNA sequencing and in situ studies of mouse atherosclerotic lesions identify *AHR* pathway expression in FMC in the media and fibrous cap

Since *AHR* shares downstream molecular pathways with *TCF21*, we investigated whether *AHR* was also a regulator of phenotypic modulation in vivo. For these experiments, we performed single-cell RNA-seq using *MYH11*-Cre^ERT^ smooth muscle cell specific lineage-tracing mouse model of atherosclerosis (SMC^*LnT/WT*^) as described previously ^12^. Briefly, we digested aortic root tissues and isolated tdTomato + cells by FACS and performed single cell RNA sequencing (scRNA-Seq). These data were analyzed with Seurat to perform a PCA-based graphical clustering, which was represented as a UMAP (Uniform Manifold and Approximation and Projection) plot (Figure 3a). We found 7 clusters within the tdTomato+ cells, and identified the markers that characterized each cluster (Supplemental Table 4). Genes that distinguished clusters 0, 3, 4, and 5 included mature SMC markers such as *Cnn1*, indicating that they represent closely related mature SMC populations (Figure 3b). On the other hand, downregulation of mature SMC markers and upregulation of genes such as *Lum* and other markers established through our previous work identified clusters 1 and 2 as representing the fibroblast-like FMC (Figure 3c).^12^ Cluster 6 expressed markers for pericytes, and the small number of cells in cluster 7 expressed markers for macrophages, which were likely contaminants. To assess activity of the *Ahr* pathway in SMC lineage cells, we surveyed expression of the prototypical downstream gene *Cyp1b1* in the single-cell analysis. We found that while *Cyp1b1* was expressed in both SMC and FMC clusters, the average expression was greater in FMC (Figure 3d, Supplemental Figure 3).

**Figure 3.**
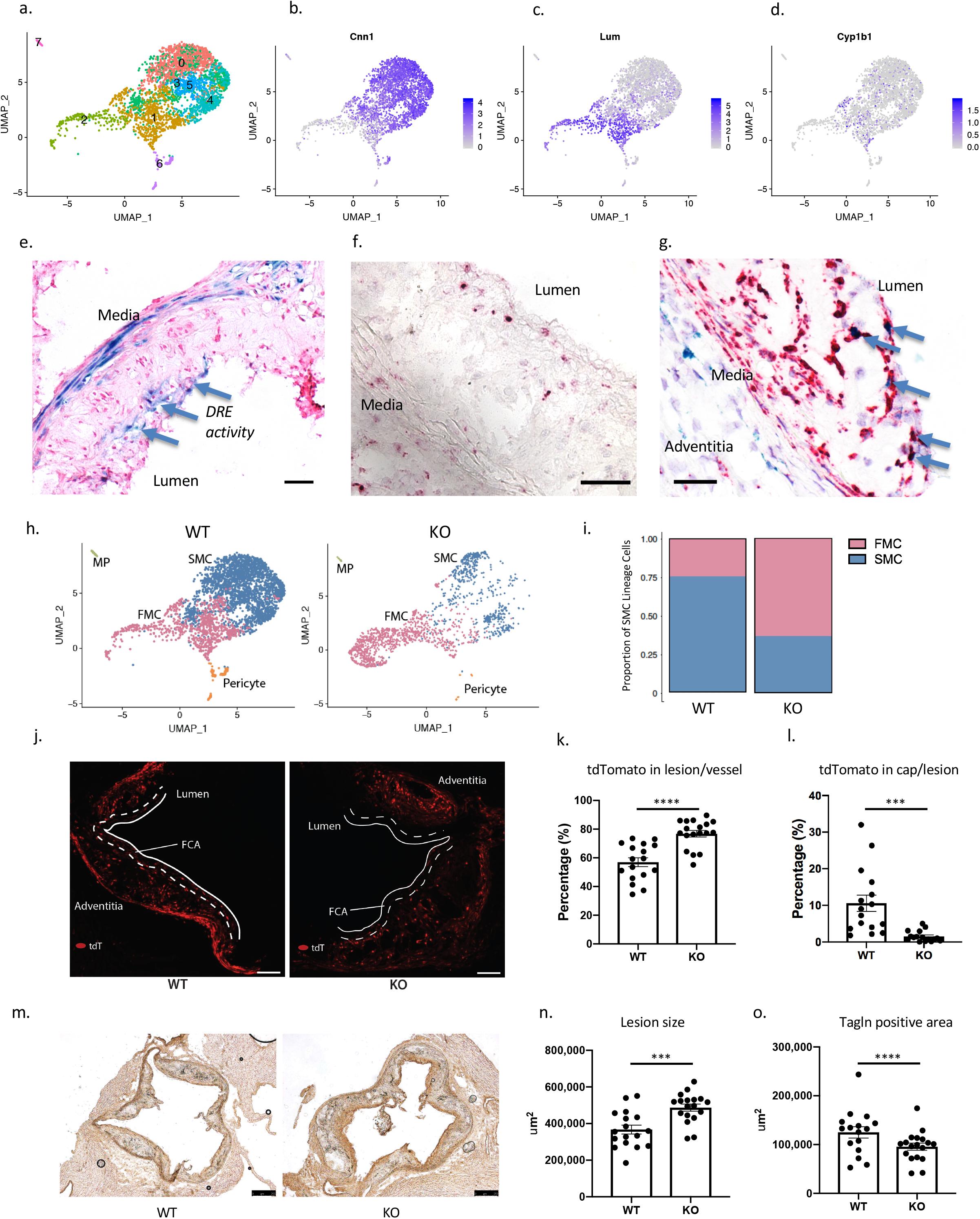
Single-cell and in situ analysis of atherosclerotic mouse aorta identifies AHR as a critical upstream regulator of phenotypic modulation and atherosclerosis. (a) UMAP plot of tdTomato+ SMC-lineage cells from SMC^*LnT/WT*^ mice (n=3327 cells). (b, c) Featureplots of SMC marker *Cnn1* and fibroblast marker *Lum* show delineation of SMC and FMC. (d) *Cyp1b1* is expressed both in FMC and SMC, but expressed at higher levels in FMC (p=1.3e-9, Supplemental Figure 4). (e) A dioxin reporter element (DRE) *lacZ* reporter transgene in ApoE null atherosclerosis model shows X-gal staining in media and the lesion cap (blue arrows) at 16 weeks of HFD (bar = 50μm). (f) RNAscope of vascular lesion of SMC^*LnT/WT*^ mouse showed expression of *Cyp1b1*(red) at the fibrous cap and in the adventitia (bar = 50μm). (g) RNAscope of lesions for *Cyp1b1* (green) and *tdTomato* (red) showed colocalization in the lesion cap (blue arrows). *Cyp1b1* staining is also seen in the media and adventitia (bar = 50μm). (h) UMAP clusters of tdTomato+ cells from WT mice (left, n=3177), assigned as SMC and FMC based on their expression of SMC and fibroblast markers. KO tdTomato+ cells (right, n=1534) show shifting of transcriptional identity towards FMC compared to WT mice. (i) There is greater proportion of FMC in KO compared to the WT (WT 25% vs. KO 63%, Chi-square p=2.7e-145). (j) The tdTomato+ signal in lesion cap of WT (left) vs. KO (right) lesion after 16 weeks of high fat diet (bar = 50μm). (k) There is an increase in the tdTomato+ area in the lesion of KO animals compared to the WT mice (p<0.0001) (l) There is a significant decrease in the tdTomato+ area within the lesion cap, defined as 30um area from the luminal surface (p=0.0004). (m) Immunohistology was performed on WT (left) and KO mice (right) for Tagln (bar = 250μm). (n, o) The lesion size was increased (p=0.0005), but the Tagln+ area was reduced in the KO (p<0.0001).

We then introduced a previously described transgene carrying the dioxin response element (DRE) *lacZ* reporter onto the *ApoE*^−/−^ model of atherosclerosis to identify the areas of the atherosclerotic lesion with *AHR* pathway activity.^14^ This transgene carries tandem repeats of the AHR binding motifs with minimal promoter driving *lacZ* expression. After 16 weeks of high fat diet, X-gal staining of the aortic sinus showed the presence of *Ahr* pathway activity in the media as well as the lesion cap (Figure 3e). We further assessed activity of the *Ahr* pathway by identifying the location of *Cyp1b1* gene expression using RNAscope in-situ hybridization. We again found that *Cyp1b1* gene expression was present in the media and the lesion cap, and that this signal in the lesion cap co-localized with tdTomato expression, confirming that the *Ahr* pathway activity in the lesion is present in the FMC population (Figures 3f, 3g).

### SMC-specific *Ahr* KO increases the proportion of modulated SMC, producing larger disease lesions with fibrous caps deficient in SMC lineage cells

In order to understand how *Ahr* affects the process of phenotypic modulation in vivo in the disease setting, we introduced *Ahr*^*-flox/flox*^ alleles onto a line that contained the tamoxifen inducible Cre expression, *Rosa26* locus *tdTomato* lineage marker, and the disease promoting *ApoE*^−/−^ alleles, producing mice with the genotype *Myh11*^*CreERT2*^, *Ahr*^*ΔSMC/ΔSMC*^, *Rosa*^*tdT/+*^, *ApoE*^−/−^ (SMC^*LnT/KO*^). Gavage with tamoxifen at 8 weeks of age generated *Ahr* knockout mice that were maintained on high fat diet for 16 weeks. At 24 weeks of age, mice were sacrificed for scRNA-Seq or histology of the aortic sinus and ascending aorta. We collected tdTomato+ cells from the aorta and conducted sequencing and analysis as described above.

We compared the SMC-specific lineage traced wild-type mice SMC^*LnT/WT*^ (WT) and the SMC^*LnT/KO*^ (KO) mice to assess the effect of *AHR* deletion on lesion characteristics. We performed scRNA-Seq on the tdTomato+ cells of WT and KO mice, and identified clustered cells as FMC, SMC, or pericytes (Figure 3h, WT left, KO right). We found that the KO mice had a significantly larger proportion of FMC compared to the WT group (WT 25% vs. KO 63%, 4711 total cells from 3 mice/group, Chi-square p=2.7e-145, Figure 3i). In order to confirm these findings in a larger cohort of mice, we compared the proportion of tdTomato+ cell populations in histology sections in 15 WT and KO mice per group (Figure 3j). There was no difference in the overall tdTomato+ area in the vessel, however, the proportion of tdTomato+ area in the intimal lesion was higher in the Ahr KO group compared to the WT group (WT 57.0±3.1% vs. KO 76.8±2.3%, p<0.0001, Figure 3k) Additionally, the tdTomato+ FMC were significantly absent from the fibrous cap area (area extending 30μm from the lumen), suggesting either failure of cells to reach the cap or their exit from the cap and ingress back into the plaque (WT 10.6±2.2% vs. KO 1.5±0.4% by area, p=0.0004, Figure 3l).

We performed Tagln immunohistology of the aortic root to characterize the atherosclerotic lesions in the WT and KO mice (Figure 3m). We found the lesion size to be increased in the KO cohort compared to the wild type (367580±24295 vs. 486949±19086 μm^2^, p=0.0005, Figure 3n). The Tagln staining revealed that the overall SMC area in the lesion was decreased in the KO mice compared to the WT (125017±11600 vs. 95748±7265 μm^2^, p<0.0001, Figure 3o), consistent with findings with the scRNA-Seq data.

### SMC-specific *Ahr* KO promotes the conversion of lineage traced SMC to a chondrogenic phenotype

In order to further characterize the phenotypically modulated SMC in the *Ahr* KO mice, we further divided the modulated population to FMC1 and FMC2 cells (Figure 4a), where FMC2 represents the population that was significantly increased in the *Ahr* KO mice compared to the WT (47.2% vs. 5.4%, chi-square p = 4.9e-262, Figure 4b). The phenotype of these cells was characterized by expression of genes such as *Col2a1*, *Spp1*, *Alpl* and *Enpp1* (Supplemental Table 4), i.e. enriched for biological pathways related to ossification, collagen fibril organization and skeletal system development (Figure 4c). We then determined the upstream regulators driving transcriptional changes (top 100 upregulated genes) between FMC2 and FMC1 using IPA (QIAGEN Inc., https://www.qiagenbioinformatics.com/products/ingenuitypathway-analysis).^15^ When these upstream regulators were queried using STRING-db,^16^ we found again that ossification and cartilage development, as well as TGF-β pathways were key biological processes responsible for promoting their phenotype and possibly regulating this transition from FMC1 to FMC2 (Figure 4d, 4e). As expected, Ahr was reported as the top down-regulated transcription factor, while the top up-regulated factors formed a closely linked network involved in bone formation, including Bmp2, Sox9, Tgfb1, Wnt3a, and Pth (Figure 4f). Based on the strong chondrocyte-like transcriptional phenotype of the FMC2 population, and by comparison to the term fibromyocyte, we termed these cells “chondromyocytes (CMC)” to reflect their SMC origin and their chondrogenic phenotype.

**Figure 4.**
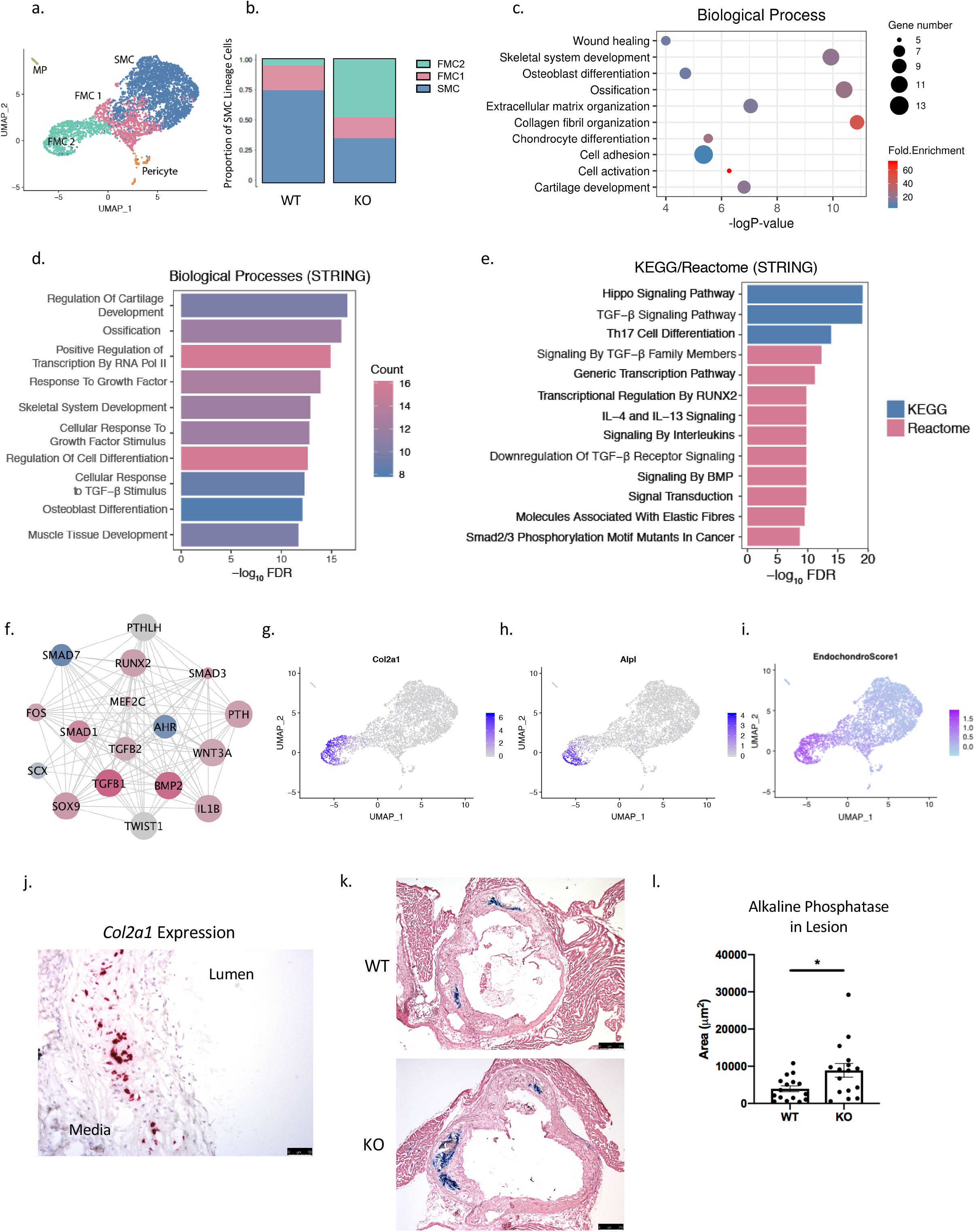
SMC-specific *Ahr* KO produces modulated cells with a chondrogenic phenotype. (a) On the combined UMAP of tdTomato+ lineage traced cells from WT and KO mice, the FMC population was further clustered to FMC1 and FMC2 population based on graph-based clustering with PCA. (b) The proportion of FMC2 cells was significantly greater for the KO compared to the WT group (Chi-square p = 4.9e-262). (c) DAVID Gene Ontology analysis of the top 100 differentially regulated genes in FMC2 compared to FMC1 population. (d) Enriched biological pathways and (e) KEGG/Reactome from the top upstream regulators of FMC1 to FMC2 transcriptional change. (f) A network of top upstream regulators of FMC2 vs. FMC1 transcriptional phenotype show AHR as top inhibited upstream factor (blue = inhibited, red = activated, size of circle = Z-score; nodes from Ingenuity Pathway Analysis, edges from STRING-db). (g) The FMC2 population was enriched for expression of chondrocyte/osteoblast markers *Col2a1* (h) and *Alpl*, and (i) an endochondral score built from the average expression of 13 genes show strong signal in the FMC2 population. (j) RNAscope of *Col2a1* expression, which is localized to the neointimal layer of the atherosclerotic lesion in mouse aortic sinus. (k) Alkaline phosphatase activity is detected in the neointima of atherosclerotic lesion at 16 weeks of HFD, in both WT and KO (bar = 250μm). (l) The average lesion area with alkaline phosphatase activity is larger in KO group compared to the WT group (3936±794 vs. 8893±1850 μm^2^, p=0.02).

### Localization and quantification of disease associated chondromyocytes

To characterize the location of CMC in transcriptomic space in relation to FMC and SMC, we visualized expression of *Col2a1*, and *Alpl* and noted that these genes identified CMC as being maximally separated from SMC, and closer to FMC, suggesting that they derive from FMC as part of a linear trajectory (Figures 4g, 4h). We then constructed an “Endochondral Score” based on the average expression of 13 representative genes from the gene ontology “Endochondral Ossification” (Extended Methods in Supplemental Material), and visualized this on a UMAP showing that the CMC population had a clear signal for endochondral ossification (Figure 4i). To define the anatomical location of CMC, we performed RNAscope as well as histochemical analysis of the atherosclerotic lesions. We found the representative CMC gene *Col2a1* to be expressed in intimal cells of the lesion but not in the lesion cap, unlike the localization that we had observed for *Cyp1b1* (Figure 4j). Alkaline phosphatase (AP) activity was also localized using a chromogenic assay in the atherosclerotic lesions of mice (Figure 4k), and AP activity was found exclusively in the intima of the lesions. Consistent with our finding from the scRNA-Seq data showing greater CMC number in the *Ahr* KO mice, we found the relative AP-stained area to be larger in the *Ahr* KO mice cohort compared to the WT (Figure 4l).

### *AHR* regulates HCASMC phenotype

We used cultured HCASMC to examine the effect of *AHR* on cell state phenotypes as predicted by the RNA-Seq and ChIP-Seq experiments. First, using a gap closure assay, we measured the effect of *AHR* knockdown on cell migration, and found an increase in the migration and cell coverage with silencing of *AHR* (p=0.048, Figure 5a). We also looked for any change in the proliferative capacity of the HCASMC with *AHR* modulation using an EdU proliferation assay. We found that AHR knockdown resulted in increased uptake of EdU, and the opposite result when *AHR* was overexpressed (p<0.0001 for both, Figure 5b). We also assayed for apoptotic activity in HCASMC subjected to apoptotic stress induced by doxorubicin. *AHR* knockdown resulted in reduced apoptotic activity based on the Annexin V assay (p=0.016, two-way ANOVA, Figure 5c). We also investigated the rate of calcification of HCASMC grown in calcification media. There was an increase in the total calcification in cells where *AHR* was silenced, and decreased calcification when *AHR* was overexpressed (Figure 5d). Overall, we found that *AHR* had an anti-migratory, anti-proliferative, anti-calcifying, and pro-apoptotic effect on HCASMC in vitro. These findings in human cells were congruent with the scRNA-Seq data in mouse.

**Figure 5.**
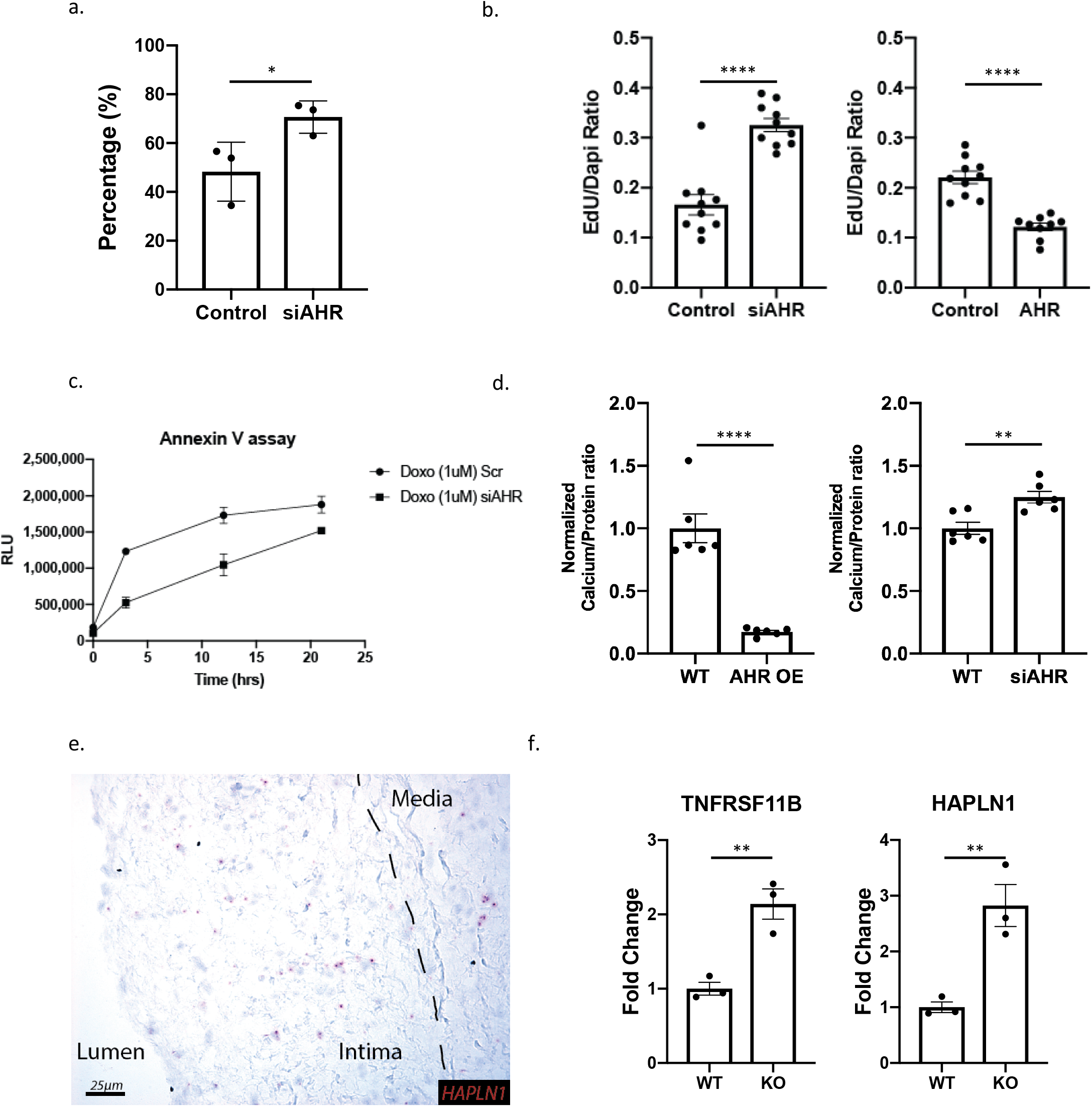
*AHR* regulates HCASMC phenotype in vitro. (a) Radial migration assay of HCASMC with scrambled control and *AHR* siRNA show increased migration with *AHR* knockdown (* p=0.049). (b) EdU uptake assay used to measure proliferation of HCASMC showed an increase with *AHR* knockdown and decrease with *AHR* overexpression (p<0.0001 for both). (c) Apoptosis quantification by Annexin V assay shows *AHR* knockdown to decrease apoptosis (p=0.016, 2-way ANOVA). (d) Calcification assay shows a decrease in calcification with *AHR* overexpression in HCASMC and an increase with AHR knockdown (p<0.0001 for *AHR* OE, p= 0.004 for *AHR* knockdown). (e) Chondrocyte markers identified in FMC2 are increased with AHR knockdown in HCASMC (p=0.0068 for *TNFRSF11B*, p=0.0094 for *HAPLN1*). (f) Chondrocyte marker *HAPLN1* is also expressed in the diseased human coronary artery as determined by RNAscope, and concentrated in the intimal layer.

### Chondromyocyte markers are expressed in human coronary artery lesion cells

We next investigated whether genes upregulated in CMC of *Ahr* KO mice were also expressed in human coronary artery cells of similar phenotype using our previously published scRNA-Seq dataset.^12^ We performed PCA clustering using Seurat, and used *MYH11* expression to identify mature contractile SMC and *LUM* to identify fibroblasts, FMC and CMC (Supplemental Figure 4a-c). Human cells expressing genes specific for the CMC population, *TNFRSF11B* and *HAPLN1*, were positioned in cluster 4 between the SMC and fibroblast clusters (Supplemental Figure 4d-g). We then applied the endochondral score on the human data, and showed that this cluster of cells had higher scores than others within the SMC-Fibroblast continuum (Supplemental Figure 4h). We probed for *HAPLN1* gene expression in the human coronary artery tissues with RNAscope, and identified staining primarily in the intima, with little staining in either the lesion cap or media (Figure 5e). Further, there was a significant increase in the expression of both markers when *AHR* was knocked down with siRNA in HCASMC (Figure 5f). Taken together, these data suggest that CMC are present in the human plaque, with CMC expressing similar markers and similarly localized in the neointima of the coronary artery plaque.

### Antisense long non-coding RNA *AC003075.4* is associated by GWAS with CAD at 7q21 where it regulates *AHR* expression

In the genomic locus at 7q.21, an LD block represented by lead SNP rs6968554 was found to be significantly associated with coronary artery disease with a look-up in the CARDIOGRAM+C4D GWAS (p=1e-4, Figure 6a). Furthermore, this variant is a *cis*-eQTL for *AC003075.4*, a lncRNA that is anti-sense to *AHR*, in the GTEX database in tibial artery (p=2.5e-4, Figure 6b). We found that knocking down *AC003075.4* resulted in the down-regulation of *AHR* expression, while overexpression resulted in the up-regulation of *AHR* expression (Figure 6c). When *AHR* was knocked down with siRNA, there was up-regulation of *AC003075.4*, and when *AHR* was overexpressed, there was down-regulation of *AC003075.4*, suggesting a negative feedback mechanism (Supplemental Figure 5). Furthermore, when *TCF21* was overexpressed, *AC003075.4* expression increased, and expression decreased with *TCF21* knockdown (Figure 6d). These finding suggest that *AHR* expression is regulated by *TCF21* and its antisense lncRNA, which are both associated with CAD phenotype by common variants.

**Figure 6.**
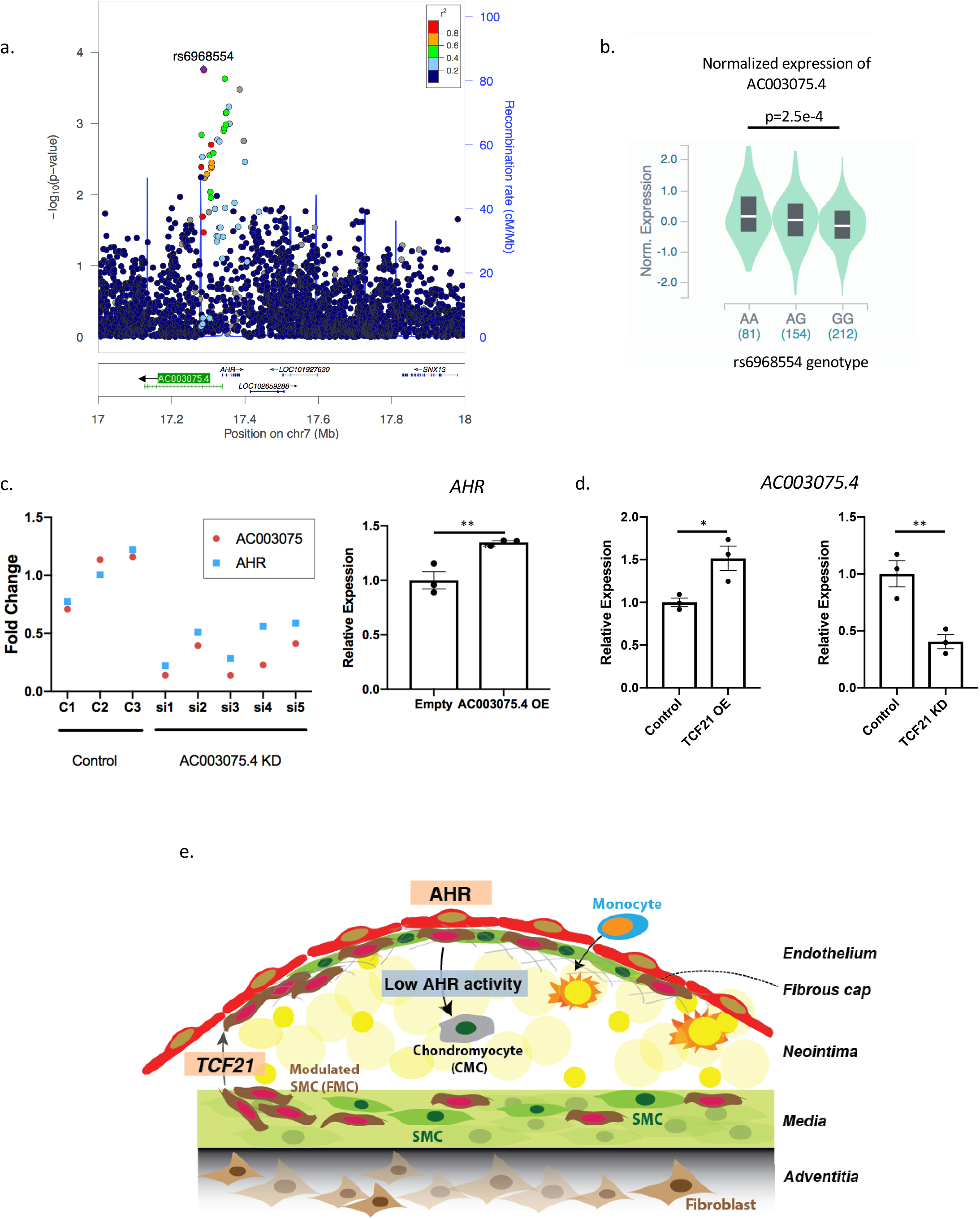
Antisense long non-coding *AC003075.4* is correlated with CAD in GWAS and regulates *AHR* expression. (a) rs6968554, a lead-SNP upstream of the *AHR* gene, is significantly correlated to CAD on CARDIOGRAM+C4D (p=1e-4). (b) In GTEx vascular tissue (tibial artery), rs6968554 is an eQTL for *AC003075.4* expression, a lncRNA that is anti-sense to *AHR (ANOVA p=2.5e-4)*. (c) *AC003075.4* and *AHR* expression are highly correlated, (*left*) *AC003075.4* knockdown decreases *AHR* expression (p = 0.0062), (*right*) AC003075.4 overexpression increases *AHR* expression (p=0.012). (d) *TCF21* positively regulates *AC003075.4* expression in HCASMC (p=0.027 for overexpression, p=0.01 for knockdown). (e) Working model of *AHR* function in the development of atherosclerosis. *TCF21* is initially up-regulated and promotes SMC to dedifferentiate and participate in lesion and fibrous cap development. *AHR* is activated in the lesion cap and plays a role in maintaining the FMC in this region, with *AHR* loss promoting FMC to migrate into the neointima and assume a chondrocyte-like phenotype, i.e., chondromyoctes (CMC).

## Discussion

Environmental factors activate gene regulatory programs to contribute to the risk for coronary artery disease (CAD) through gene-environment interactions, and we have investigated the aryl hydrocarbon receptor (*AHR*) as a critical mediator of such effects. *AHR* is an inducible transcription factor that was first recognized for its ability to mediate the effect of a number of xenobiotic ligands such dioxin and PAH, which are established components of tobacco smoke, and its functional role in this regard considered harmful.^4,17^ However, *AHR* is now recognized as an important developmental factor and physiological regulator in the vascular, immune and reproductive systems.^6,18,19^ *AHR* regulates cell cycle and cellular differentiation through interaction with numerous other signaling pathways, including those regulated by TGF-β and Notch.^20,21^ Many of these actions are likely mediated by endogenous ligands, and for the most part considered beneficial. Thus, experiments described in this work have focused on the function of *AHR* in vascular smooth muscle cells in vascular disease, in the absence of known external environmental ligand activation.

*AHR* was identified as a downstream target of the CAD associated factor *TCF21*, which we have recently shown to promote phenotypic modulation as a mechanism of risk prevention.^9,12^ In studies reported here, we gained further insight into the regulatory interaction between *AHR* and *TCF21*, and found a significant co-localization of AHR and TCF21 binding in the HCASMC genome in regions of open chromatin. Furthermore, we found a significant overlap between the transcriptional changes induced by *AHR* and *TCF21* modulation that point to critical pathways involved in SMC modulation, including cell migration, differentiation, and cell cycle. Importantly, *AHR* showed a unique gene expression signature suggesting inhibition of ossification. And while *AHR* was found to regulate genes that modulate SMC migration and proliferation, the direction of effect as identified by in vitro assays was opposite to that of *TCF21*.^10^ These data thus suggested a distinct functional profile compared to *TCF21*.

We have previously used lineage tracing and scRNA-Seq to show that *Tcf21* promotes the modulation of SMC to fibroblast-like cells that we termed fibromyocytes (FMC),^12^ and were thus interested to determine how *Ahr* might affect SMC phenotype in the disease setting. In contrast to the reduced FMC in lesion and lesion cap seen in the SMC-specific knockout of *Tcf21*, we found that there was an increase in SMC modulation with SMC-specific knockout of *Ahr*. Interestingly, loss of *Ahr* was found to promote further modulation of FMC to a chondrocyte-like cell phenotype. This is consistent with findings in model systems where *AHR* has been shown to suppress expression of pro-chondrocyte genes *Sox9*, and *Runx2*.^22,23^ Also, by analogy to the de-differentiated FMC, previous reports suggest that *AHR* may maintain the “stemness” of progenitor cells. *Watson et al.* reported that AHR mediates inhibition of osteoblast differentiation in human mesenchymal stem cells^24^, and another recent report found that *AHR* pathway activation by an endogenous ligand can maintain the undifferentiated state of embryonic stem cells.^25^ Hence, the FMC may in fact be a form of multi-potent cell population that is derived from de-differentiated SMC population, ready to differentiate to a mesenchymal lineage cell-type, including the CMC and also back to a differentiated, quiescent SMC population. The chondrogenic cells described here were previously identified as the most de-differentiated portion of the FMC, but not characterized specifically as a discrete cell lineage.^12^ Given evidence from these studies showing that these cells are independently regulated with respect to cellular phenotype, we have elected to recognize them as a discrete group of modulated SMC, and use the term chondromyocyte to distinguish them from their mature SMC origin, and their related and presumably intermediate phenotypic FMC precursors.

Thus, findings reported here contribute to our understanding of the SMC response to stress in the setting of vascular disease, but suggest that the process of phenotypic modulation is more complex than previously appreciated.^12,26^ Although it is not currently possible to integrate all available information, we propose the following model (Figure 6e). *TCF21* expression is activated early in the disease process to promote SMC de-differentiation, proliferation and migration out of the media. Subsequently, *AHR* is upregulated in FMC and serves to maintain the FMC phenotype as these cells migrate to the fibrous cap. As seen in the atherosclerotic lesions of DRE-*LacZ* reporter mouse as well as the *Cyp1b1* RNAscope performed on lesions of SMC lineage tracing mice, the *AHR* pathway becomes significantly activated in the fibrous cap. Loss of *AHR* results in FMC with greater migratory capacity reaching the cap and then moving into the plaque where they adopt a chrondrogenic phenotype, consistent with a recently proposed mechanism of cellular disease construction,^27^ or simply adopting a chondrogenic phenotype and expanding in the neointima where they contribute to lesion size and detract from cells contributing to the fibrous cap. Undoubtedly, additional genes such as KLF4 and OCT4 also control various aspects of the phenotypic modulation process,^26,28^ and their roles will be better understood as scRNA-Seq data becomes available for knockout models of these and other relevant genes.

Given the upregulation of *AHR* by *TCF21* and the overlap of regulated genes downstream, we expected that it would have a similar function to *TCF21* in SMC. However, we found that its functional profile with in vitro assays was distinct to *TCF21*, and that its loss led to an increase in the number of modulated SMC. While this profile would be expected to decrease disease risk based on our previous studies of *TCF21*, the phenotype of the increased SMC-derived modulated cells in the knockout mice was quite different from the FMC we characterized previously, and more consistent with a chondrogenic type cell. Further, the causal relationship of the human data is complex, but suggests a protective role of *AHR* as the major disease allele “G” for lead SNP rs6968554 is associated with lower expression of regulatory lncRNA *AC003075.4* and thus *AHR*. However, additional studies are required to better understand the complex relationship between the *AHR* gene, SMC phenotypic modulation associated with variance of this gene, and the directionality of disease risk.

## Conclusion

In this study, we provide evidence that the *AHR* pathway, mediated by an environment-sensing transcription factor, is active in modulated SMC, and can inhibit the phenotypic modulation of SMC to chondrocyte-like cells during development of atherosclerosis.

## Methods

### Culture of human coronary artery smooth muscle cell (HCASMC)

Primary human coronary artery smooth muscle cells (HCASMC) were purchased from Cell Applications, Inc (San Diego, CA) and were cultured in complete smooth muscle basal media (Lonza, #CC-3182) according to the manufacturer’s instructions. All experiments were performed with HCASMC between passages 5-8. HEK293 cells were maintained in DMEM containing high glucose, sodium pyruvate and L-glutamine supplemented with 10% FBS.

### Knockdown and overexpression

For the siRNA transfection, cells were grown to 60% confluence, then treated with siRNA or scramble control to final concentration of 20nM with RNAiMax (Invitrogen, Carlsbad, CA) for 12 hours. The siRNA for *AHR* and *TCF21* were purchased from Origene (SR300136, SR321985), and siRNA for *AC003075.4* was custom designed (Ambion/Life Technologies Corp, Supplemental Table 1). The cells were collected 48 hours after transduction. For the overexpression study, HCASMC were transduced with 2nd generation lentivirus with *AHR* cDNA (HG10456-CY, Sino Biological) and *AC003075.4* cDNA (custom gBlock fragment, IDT) cloned into pWPI (Addgene #12254) using NEBuilder HiFi cloning (New England Biolabs). Briefly, for lentiviral transduction, the cells were treated at 60% confluence with virus at MOI of 5 for 24 hours. The virus was removed and replaced with low-serum media for 48 hours prior to collection for downstream applications.

### RNA isolation and qRT-PCR

RNA was isolated using RNeasy mini kit (Qiagen) and total cDNA was prepared using iScript cDNA synthesis kit (Biorad, Hercules, CA). Gene expression levels were measured using Taqman probes (Invitrogen) (*AHR, TNFRSF11B, HAPLN1*) and custom probes (*AC003075.4*, see *supplemental material*) for SYBR Green assay and quantified on a ViiA7 Real-Time PCR system (Applied Biosystems, Foster City, CA) and normalized to *GAPDH* levels.

### RNA-sequencing

Four replicates were used for each sample. The RNA were processed and analyzed as described previously. Briefly, RNAs were sent to Novogene for sample QC, library preparation, and sequencing. All samples passed QC, and 250-300 bp insert cDNA libraries were prepared for each sample. Subsequently, sequencing was performed on a Novaseq 6000 platform with paired-end 150 bp reads. The script for RNA-Seq analysis can be found in github (https://github.com/milospjanic/rnaSeqFPro). Additionally, the web-based tools iDEP.90 (http://bioinformatics.sdstate.edu/idep/) was used for analysis of the RNA-Seq data and visualization using the counts data generated from FeatureCounts.^29^

### ChIP assay and ChIP-Seq

ChIP was performed using AHR antibody (Santa Cruz, sc-5579) and TCF21 antibody (HPA013189, Sigma) according to previous published protocols.^30^ The purified DNA was assayed with qPCR for CYP1B1 DRE region binding to confirm successful immunoprecipitation (Supplemental Table 5).

Library for ChIP-Seq was prepared using standard procedures as decribed previously.^30^ The script for RNA-Seq analysis can be found in github (https://github.com/zhaoshuoxp/Pipelines-Wrappers/blob/master/ChIPseq.sh). *macs2 bdgdiff* was used for differential peaks calling with default parameters. We utilized the Genomic Regions Enrichment of Annotations Tool (GREAT 3.0) to analyze the detected peaks for Gene Ontology. The *HOMER findMotifsGenome*.*pl* script was employed to search for known TRANSFAC motifs and to generate de novo motifs. The *intersectBed* was used to find at least 1 bp overlapped peaks between AHR and TCF21. *P* values were calculated by *Fisher’s exact test with the whole genome as the background*.

### ATAC-Seq

For ATAC-Seq, we followed the published Omni-ATAC-seq protocol.^31^ For qPCR experiments, the purified DNA was quantified with ABI ViiA 7 and Power SYBR Green Master Mix (ABI 4368706) and normalized by genomic DNA, which was extracted using Quick-DNA Microprep Kit (Zymo D3020). Assays were repeated at least three times. Data shown were average values ± SE of representative experiments. Libraries were sequenced on HiSeq X10 for 150-bp paired-end sequencing and analyzed as described previously.^30^ The script for RNA-Seq analysis can be found in github (https://github.com/zhaoshuoxp/Pipelines-Wrappers/blob/master/ATACseq.sh).

### Co-immunoprecipitation (co-IP) and ChIP-Western Blot

HEK293 cells were transfected with plasmids carrying HA-tagged AHR cDNA (HG10456-CY, Sino Biological) and MYC-tagged *TCF21* cDNA (HG16907-CM, Sino Biological). The cells were then treated with 10nM TCDD or control (10nM DMSO) for 1 hour. For the co-IP experiment, lysates were collected using RIPA buffer, and assay was performed with Nuclear Complex Co-IP Kit (Active Motif) as instructed using antibodies against HA-tag (Thermo Fisher cat#26183), MYC-tag (Abcam, cat#ab32) or IgG at 1:5000 concentrations. The immunoprecipitate was blotted against the second protein (MYC or HA) for visualization. For the ChIP-WB, the treated HEK293 cells were prepared using protocol for ChIP, and IP was performed using antibody against HA and MYC. The immunoprecipitated product was blotted against MYC or HA antibodies.

### Mouse Models

The DRE-LacZ strain was re-derived from cryopreserved embryos at the Jackson Laboratories (*B6.Cg-Tg(DRE-lacZ)2Gswz*/J strain, JAX# 006229).^14^ The mice were genotyped for presence of LacZ, then bred to ApoE^−/−^ background.

To enact SMC-specific lineage tracing and *Ahr* knockout, we used mice containing a well-characterized BAC transgene that expresses a tamoxifen-inducible Cre recombinase driven by the SMC-specific *Myh11* promoter (*Tg*^Myh11-CreERT2^, JAX# 019079)^26,32,33^. These mice were bred with a floxed tandem dimer tomato (tdT) fluorescent reporter line (B6.Cg-*Gt(ROSA)26Sor*^*tm14(CAGtdTomato)Hze*^/J, JAX# 007914) to allow SMC-specific lineage tracing. *Ahr*^*flox/flox*^ mice were obtained from Jackson Labs (JAX#006203). All mice were bred onto the C56BL/6, *ApoE*^−/−^ background. Final genotypes of SMC lineage-tracing (SMC^*LnT/WT*^) mice were: *Tg*^*Myh11-CreERT2*^, *Ahr*^+/+^, *ROSA*^*tdT/tdT*^, *ApoE*^−/−^. Final genotypes of SMC lineage-tracing, Ahr knockout (SMC^*LnT/KO*^) mice were: *Tg*^*Myh11-CreERT2*^, *Ahr*^*ΔSMC/ΔSMC*^, *ROSA*^*tdT/tdT*^, *ApoE*^−/−^. As the Cre-expressing BAC was integrated into the Y chromosome, all lineage tracing mice in the study were male. The animal study protocol was approved by the Administrative Panel on Laboratory Animal Care (APLAC) at Stanford University. Two doses of tamoxifen, at 0.2mg/gm bodyweight, were administered by oral gavage at 8 weeks of age, with each dose separated by 48 hours to induce the lineage marker and *Ahr* knockout. Approximately 48 hours after the second dose of tamoxifen, high fat diet (HFD) was started (Dyets #101511, 21% anhydrous milk fat, 19% casein, 0.15% cholesterol). The Cre-mediated excision was confirmed in the DNA obtained from vascular tissue (Supplemental Figure 6, Supplemental Table 5 for primers).

### Single-cell RNA-Seq

For both SMC^*LnT/WT*^ and SMC^*LnT/KO*^ genotype, three mice were used at 16 weeks of high fat diet. Mouse aortic root dissociation and processing of cells was described previously.^12^ Briefly, dissociated cells were sorted for live/dead signal and tdTomato^+^ signal. tdTomato+ cells (considered to be of SMC lineage) were then captured for all subsequent analyses. All single-cell capture and library preparation was performed at the Stanford Functional Genomics Facility (SFGF) using 10X Genomics platform, and sequenced on an Illumina HiSeq4000 instrument.

Fastq files from each experimental group were aligned to the reference genome using CellRanger software (10X Genomics). The dataset was then analyzed using the R package *Seurat* as described previously.^34^ Briefly, the gene expression values underwent library size normalization using the published *sctransform* function.^35^ Principal component analysis was used for dimensionality reduction, followed by clustering in PCA space using a graph-based clustering approach. UMAP was then used to visualize the resulting clusters in two-dimensional space. Fastq files and matrices from the single-cell RNA-Seq experiments will be deposited in the GEO database. Primary accession codes are pending.

### Immunohistochemistry

The mouse aortic root was fixed in 0.4% PFA and embedded in OCT, and slides were prepared and processed as described previously.^12^ Sections were incubated overnight at 4C with an anti-SM22alpha rabbit polyclonal primary antibody (Abcam #ab14106, 1:300 dilution). For the alkaline phosphatase enzymatic assay, Ferangi Blue Chromogen kit (SKU#FB813H, Biocare Medical) was used as instructed. The processed sections were visualized using Leica DM5500 microscope and images were obtained using Leica Application Suite X software. Areas of interest were quantified using *ImageJ* (NIH) software, and compared using a two-sided t-test. The lesion cap was defined as 30μm segment from the luminal surface as previously described.^12,26^ Researchers were blinded to the genotype of the animals until completion of the analysis.

### HCASMC Phenotypic Assays

The Radius Cell Migration Assay kit (Cell Biolabs, Inc) was used for the gap closure assay. Briefly, HCASMC was plated at 80% confluence, then following the protocol, the polymer at the center of each well were dissolved to allow migration. Migration was quantified by remaining area after migration, compared to original area. For the proliferation assay, EdU was introduced into the cell culture 3 hours before assay for uptake. The protocol for Click-iT Plus EdU proliferation kit (Thermo Fisher) was followed as instructed. For the apoptosis assay, HCASMC was treated with Doxorubicin (1uM) for 24 hours to induce apoptosis. The RealTime-Glo Annexin V Apoptosis kit (Promega) was used as instructed to quantify the degree of apoptosis in HCASMC. To assay for calcification, HCASMC was exposed to calcification media with 10mM beta-glycerophophate and 100ug/ml ascorbic acid as described previously.^36^ The HCASMC were treated with siRNA and lentivirus for AHR as described in “Knockdown and overexpression” and then exposed to calcification media with 1% FBS for 10 days. The calcified cells were treated with 0.6N hydrochloric acid for 24 hours, then the supernatant was collected for calcium assay, and the cell layer was collected for protein quantification. The Calcium Colorimetric Assay Kit (Sigma-Aldrich) was used for quantification of calcium, then normalized to total protein content.

### Statistics

R or GraphPad Prism 7.0 was used for statistical analysis. For motif and gene enrichment analyses, we used the *cumulative binomial distribution* test. For overlapping of genomic regions or gene sets, we used *Fisher’s exac*t test and/or the *hypergeometric* test, as indicated. For comparisons between two groups of equal variance, an unpaired two-tailed Student’s *t-test* was performed. For multiple comparison testing, one-way analysis of variance (ANOVA) accompanied by Tukey’s *post hoc* test were used as appropriate. All error bars represent standard error of the mean (SE). Number of stars for the *P* values in the graphs: \*\*\*\**P* < 0.0001, \*\*\**P* < 0.001, \*\**P* < 0.01, \**P* < 0.05.

