## Supplementary Figures for "The environment-sensing aryl-hydrocarbon receptor inhibits the chondrogenic fate of modulated smooth muscle cells in atherosclerotic lesions"

Supplemental Figure 1

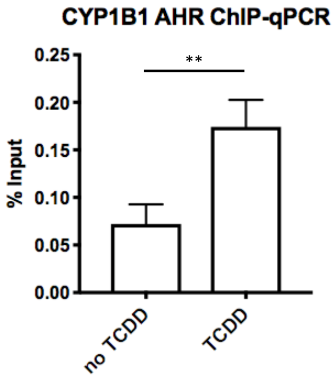

Supplemental Figure 2.

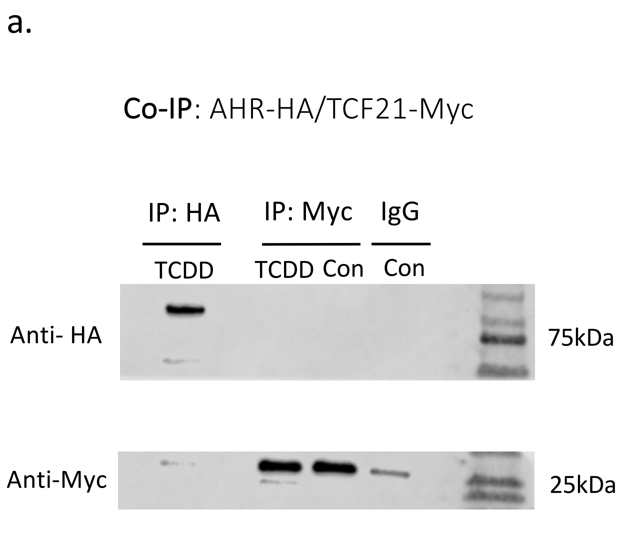

Supplemental Figure 3.

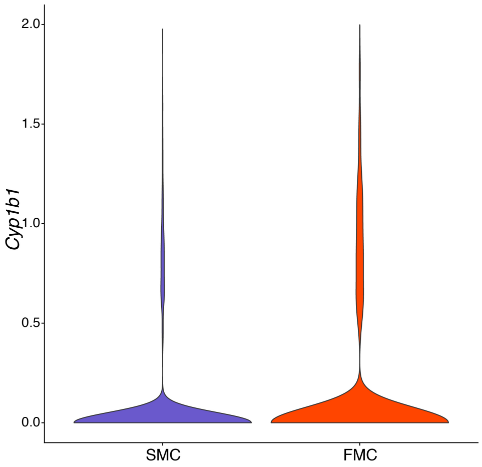

Supplemental Figure 4

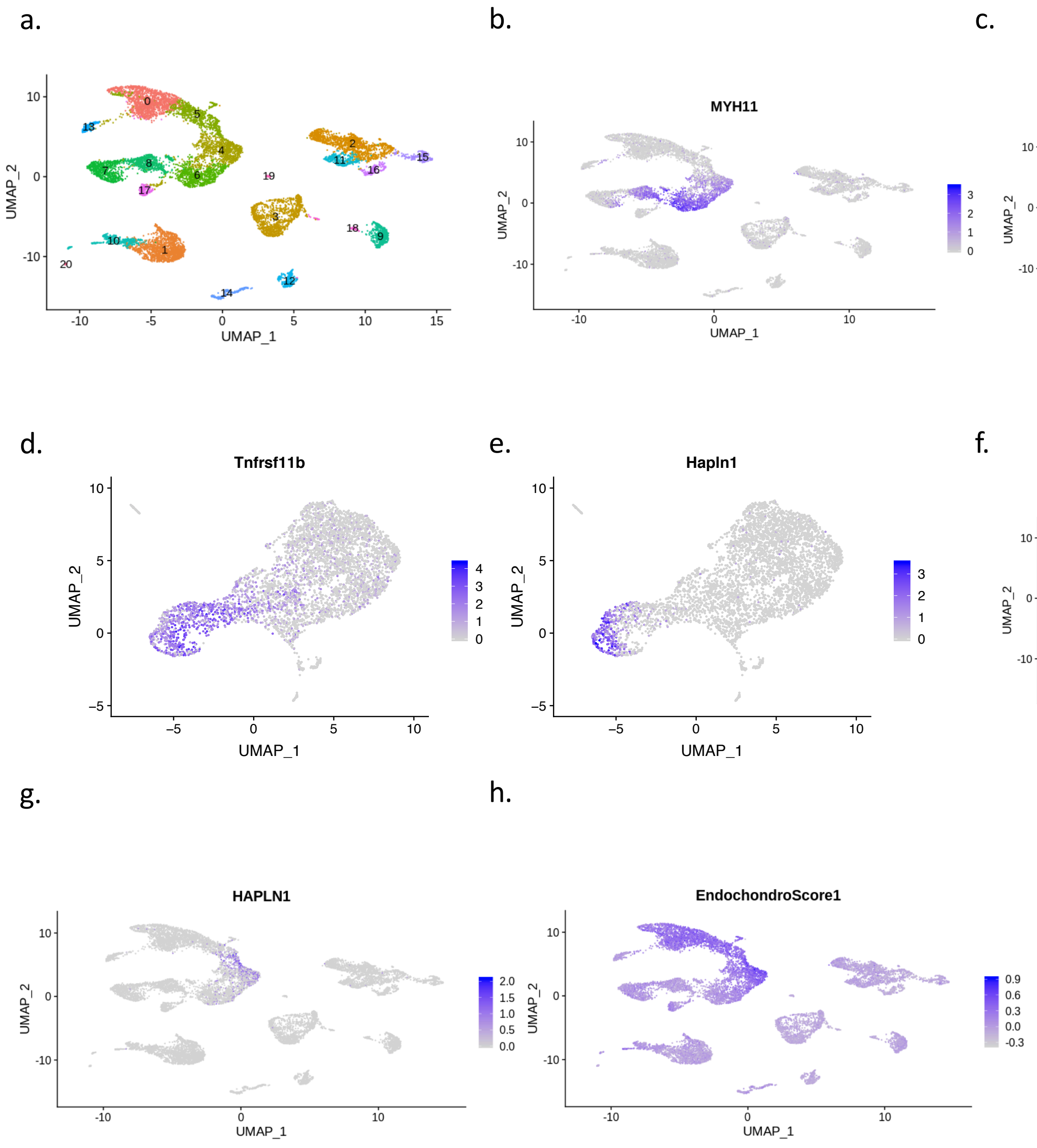

Supplemental Figure 5.

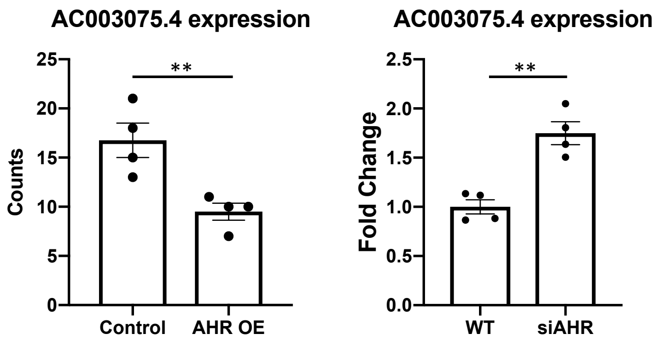

Supplemental Figure 6

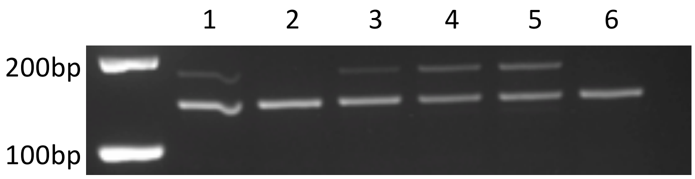
